# Movement context modulates neuronal activity in motor and limbic-associative domains of the human subthalamic nucleus

**DOI:** 10.1101/392936

**Authors:** Odeya Marmor, Pnina Rappel, Dan Valsky, Atira S Bick, David Arkadir, Eduard Linetzky, Or Peled, Idit Tamir, Hagai Bergman, Zvi Israel, Renana Eitan

**Affiliations:** Department of Medical Neurobiology (Physiology), Institute of Medical Research – Israel-Canada, the Hebrew University-Hadassah Medical School, Jerusalem, Israel; The Edmond and Lily Safra Center for Brain Research, the Hebrew University, Jerusalem, Israel; The Brain Division, Hadassah-Hebrew University Medical Center, Jerusalem, Israel; The Center for Functional and Restorative Neurosurgery, Hadassah-Hebrew University Medical Center, Jerusalem, Israel; The Functional neurosurgery program, Department of Neurological Surgery, University of California San Francisco, San Francisco, AC, USA; Functional Neuroimaging Laboratory, Brigham and Women’s Hospital, Department of Psychiatry, Harvard Medical School, Boston, MA, USA

**Keywords:** Movement inhibition, Movement planning, Subthalamic nucleus, Multiunit activity, Parkinson’s disease, Deep brain stimulation

## Abstract

To better understand the mechanism of movement facilitation and inhibition in the subthalamic nucleus (STN), we recorded subthalamic multiunit activity intra-operatively while parkinsonian patients (n=43 patients, 173 sites) performed increasingly complex oddball paradigms: auditory (‘None-Go’, n=7, 28), simple movement (‘All-Go’, n=7, 26) and movement inhibition (‘Go-NoGo’, n=29, 119) tasks. To enable physiological sampling of the different subthalamic nucleus domains in both hemispheres, each patient performed one of the oddball paradigms several times.

The human STN responded mainly to movement-involving tasks: movement execution at the motor STN and movement planning at the limbic-associative STN. In the limbic-associative STN, responses to the inhibitory cue (deviant tone) in the movement inhibition task were not significantly different from the simple movement task. However, responses to the go cue (frequent tone) were significantly reduced. The reduction was mainly in the negative component of the evoked response amplitude. Successful movement inhibition was correlated with higher baseline activity before the inhibitory cue.

We suggest that the STN adapts to movement inhibition context by selectively decreasing the amplitude of neuronal activity. Thus, the STN enables movement inhibition not by increasing responses to the inhibitory cue but by reducing responses to the release cue. The negative component of the evoked response probably facilitates movement and a higher baseline activity enables successful inhibition of movement. These discharge modulations were found in the ventromedial, non-motor domain of the STN and therefore suggest a significant role of the associative-limbic domains in movement planning and in global movement regulation.

## Introduction

Neural circuits of response inhibition are associated with the Subthalamic nucleus (STN). Classical models of the basal ganglia (DeLong, 1990, Mink, 1996), describe an excitatory input from the cortex to the STN via the hyper-direct pathway (Nambu et al., 2002). The STN exerts an excitatory influence on the output nuclei of the basal ganglia which in turn inhibit the thalamus and the cortex. Activation of the STN during inhibition of movement has been found in many fMRI, local field potential (LFP) and single unit studies in both animals and humans (Aron et al., 2006, Forstmann et al., 2012, Ray et al., 2012, Alegre et al., 2013, Schmidt et al., 2013, Bastin et al., 2014, Rae et al., 2015, Fischer et al., 2017a). The STN is also involved in response inhibition of non-motor modalities such as working memory and decision making (Brittain et al., 2012, Zaghloul et al., 2012, Wessel et al., 2016b, Herz et al., 2018).

In addition to the role of the STN in movement inhibition, the STN is also involved in the planning and execution of movement (Thobois et al., 2000, Cassidy et al., 2002, Levy et al., 2002, Foffani et al., 2004, Kuhn et al., 2006, Androulidakis et al., 2007, Oswal et al., 2013, Fischer et al., 2017b). Part of the movement planning process involves the global regulation of readiness for movement which depends on the context of the movement. In the cortex for example, a subconscious readiness potential precedes the time of voluntary movement and regulates movement execution or inhibition (Libet et al., 1983, Keller et al., 1990, Schultze-Kraft et al., 2016).

The role of the STN in global movement regulation has not been well explored although its physiology makes it eminently suitable for this function. The STN has a high spontaneous firing rate (Wichmann et al., 1994) and it tonically inhibits the thalamus via the output nuclei of the basal ganglia. Thus, the STN could be involved in the regulation of global readiness for movement.

Many studies have examined STN-mediated motor inhibition using classical versions of stop signal tasks (Kuhn et al., 2004, Aron and Poldrack, 2006, Alegre et al., 2013, Schmidt et al., 2013, Benis et al., 2016). These studies compared no-go and go trials or successful and unsuccessful stop trials. Although several studies have compared different levels of anticipation to the inhibitory signal (Ray et al., 2012, Benis et al., 2014, Fischer et al., 2016), STN-mediated mechanisms of readiness for movement in the context of motor execution or inhibition has not been well studied.

In this study we compared human STN multiunit activity on oddball tasks with three levels of movement: first, passive listening (‘None-Go’ task: no movement); second, adding presses to all tones (‘All-Go’ task: movement with a facilitation context); and third, adding inhibition of movement after the deviant tones (‘Go-NoGo’ task: movement with inhibition context). We used a similar auditory paradigm in all tasks to control for the auditory passive listening process and to directly compare simple motor planning (the All-Go task) to inhibitory motor planning (the Go-NoGo task). This enabled the investigation of the role of the STN in the execution of movement in the context of no movement, movement facilitation and movement inhibition.

## Methods

### Patients

Parkinson’s disease patients (n=43) undergoing STN DBS took part in this study. All patients met the accepted inclusion criteria for DBS surgery and gave their written informed consent. This study was authorized and supervised by the IRB of Hadassah Medical Center (reference code: 0168-10-HMO). All recordings were performed while the patients were awake and off medications (over-night washout).

### Study paradigm

We used three tasks as illustrated in Fig. 1, A. In all three tasks, series of 120 tones with two different pitches were played in a pseudo-random order. The frequent tones (82% of the played tones) were delivered at a high pitch (1200Hz) and the deviant tones (18% of the played tones) were delivered at a low pitch (300Hz). The tone duration was 250ms followed by a 1000ms pause (the total inter- trial interval was 1250ms) and the total duration of the task was 2.5 minutes on all three tasks. The difference between the tasks was the instruction to the participants: 1. ‘None-Go’ task: Participants were not informed about the task; i.e., the tones were played without any instruction to the participants (but we verified that the patient was awake). 2. ‘All-Go’ task: Participants were instructed to press a hand button as fast as possible after each tone (both frequent and deviant tones). 3. ‘Go-NoGo’ task: Participants were instructed to press a hand button as fast as possible after the frequent tones and not to press the button after the deviant tones.

**Figure 1.**
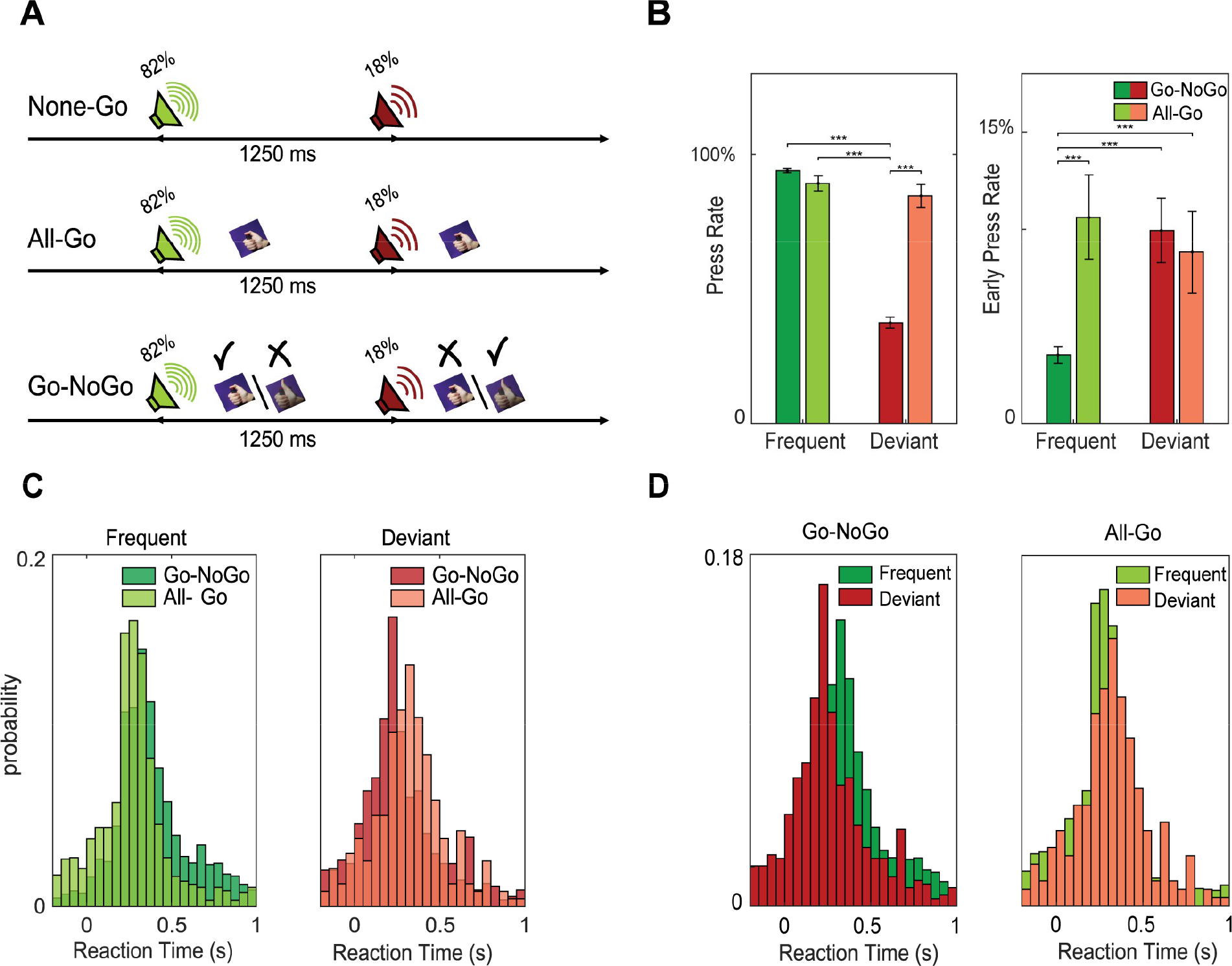
Task Paradigm and Behavioral Results. **A.** Task paradigm: None-Go – The subject listened passively to a played oddball paradigm. All-Go - The subject was also instructed to press a handed button as fast as possible after each tone of the played oddball paradigm. Go-NoGo - The subject was instructed to press the button as soon as possible only after the frequent tone, and not to press after the deviant tone. **B.** Press rate and early press rate to the frequent (green) and deviant (red) tone in the All-Go (light) and Go-NoGo (dark) tasks. Early press was defined as a press that was 0-200ms before tone onset. **C.** Distribution of reaction times in the All-Go (light green/red) and Go-NoGo (dark green/red) tasks to the frequent (left) and deviant (right) tones. **D.** Distribution of the reaction times of the frequent and deviant tones in the Go-NoGo and All-Go tasks.

Neuronal data were recorded in different areas along the left or right dorsolateral-ventromedial STN axis (see details below) while the participants performed the tasks. Recordings of all three tasks (None-Go, All-Go and Go-NoGo) in all four recording sites (right dorsolateral STN, left dorsolateral STN, right ventromedial STN, left ventromedial STN) would have been preferable. However, recordings during surgical navigation is limited by the additional clinical risks for the patient as well as by the patient’s attention span. Prior to each recording session, the neurosurgeon (ZI) verified the clinical state of the patients (for example, no excessive increased cerebrospinal fluid leak) and approved carrying out the recording session. To further minimize the clinical risk, the total recording time for research purposes for each patient was a maximum of 8 minutes. Therefore, each patient was engaged in only one task, which was repeated in the four STN recording domains. Tones and press times were saved with neuronal data on the same data acquisition device (MicroGuide or NeuroOmega, AlphaOmega, Nazareth, Israel). All patients reported right hand dominance. Participants were asked to press the button using their right thumb or index finger while recordings in the left (contra-lateral) or right (ipsi-lateral) STN. Since the Go-NoGo task instructions were thought to be too complex for some of the patients, the Go-NoGo group of patients were trained on the task prior to surgery.

### Surgery

The surgical technique is described elsewhere (Zaidel et al., 2009). Briefly, surgery was performed using the CRW stereotactic frame (Radionics, Burlington, MA, USA). STN target coordinates were chosen as a composite of indirect targeting based on the anterior commissure-posterior commissure atlas-based location, and direct targeting with three Tesla T2 magnetic resonance imaging (MRI), using Framelink 5 or Cranial software (Medtronic, Minneapolis, USA). A typical trajectory was ~60° from the axial anterior commissure-posterior commissure plane and ~20° from the mid-sagittal plane. Final trajectory plans were slightly modified to avoid the cortical sulci, ventricles and blood vessels (as seen in T1 scans with contrast media).

### Electrophysiological recordings

The microelectrode recording data were acquired with the MicroGuide or the NeuroOmega systems (n= 19 and 24 patients respectively, AlphaOmega Engineering, Nazareth, Israel) as previously described (Marmor et al., 2017). Neurophysiological activity was recorded via polyamide coated tungsten microelectrodes with an impedance of approximately 0.5 MΩ (measured at 1000Hz). For the MicroGuide system, the signal was amplified by 10,000, band-passed filtered from 250 to 6000 Hz using a hardware four-pole Butterworth filter, and sampled at 48 kHz by a 12-bit A/D converter (using ±5 V input range). For the NeuroOmega system, the signal was amplified by 20, band-passed filtered from 300 to 9000 Hz using a hardware four-pole Butterworth filter, and sampled at 44 kHz by a 16-bit A/D converter (using ±1.25 V input range).

Typically, two parallel electrodes separated by 2mm for each STN trajectory were advanced simultaneously along the planned trajectory. Recording began 10 mm above the presumed target (estimated by the pre-operative imaging). Electrodes were advanced into the STN in discrete steps of ~0.1 mm. The task was performed several times (2.4±1.2) along the tract in the STN while maintaining the electrodes stationary.

Detection of the STN entry and exit as well as differentiating between the dorsolateral oscillatory region (DLOR, sensorimotor domain) and ventromedial non-oscillatory region (VMNR, limbic-associative domain) of the STN were automatically delimited by a hidden Markov model (HMM, Zaidel et al., 2009). Recording locations in the STN subdomains are presented in Fig. 2A-C. Each recording site was classified according to the automatic HMM algorithm and verified / corrected by an experienced electrophysiologist (OM). Only locations that could be defined with certainty within the STN were included in the analysis (173 out of 196 recording sites). Total STN axis length was 4.6±2.0 mm and 4.8±2.1 mm for right and left STN, respectively. STN DLOR axis length was 2.0±1.6 and 2.1±1.6 for right and left STN, respectively.

**Figure 2.**
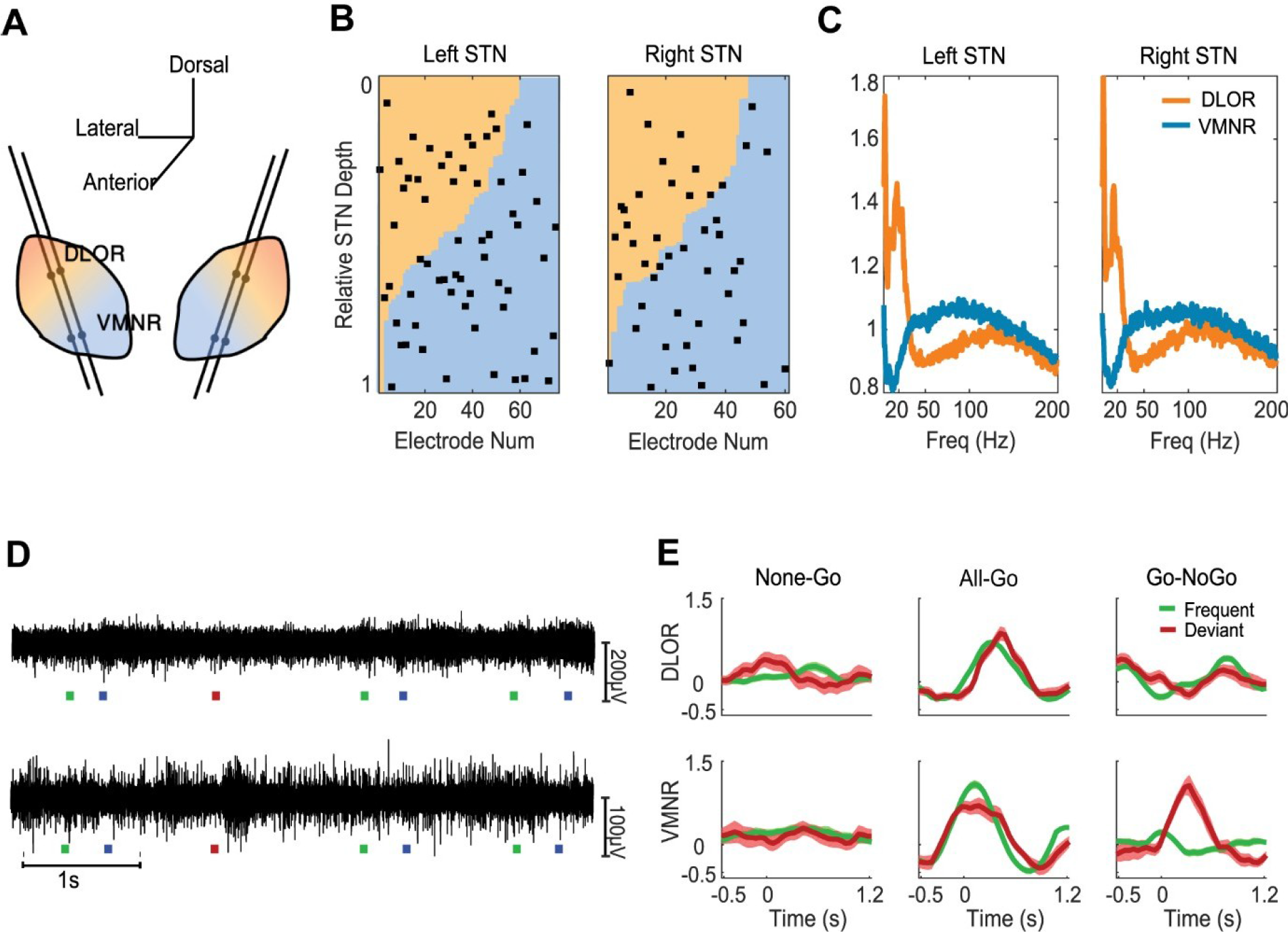
Recording Methods. **A.** Schema of the recording trajectories locations in the STN. **B.** Distribution of recording locations along the STN trajectories. Recording locations are presented by decreasing DLOR proportion and normalized by STN length. DLOR and STN lengths were automatically detected by a hidden Markov model (HMM) algorithm. **C.** The mean power spectral density (PSD) of all recording depths that were categorized as DLOR (orange) and VMNR (light blue). D&E show recording sites and trajectories in the Go-NoGo task. The recording sites were similarly distributed in the All-Go and None-Go tasks (data not shown). **D.** Examples of 5 second microelectrode high filtered signals (300-9000Hz) during the Go-NoGo task. Colored squares at the bottom mark the times of the frequent tone (green), deviant tone (red) and presses (blue). Upper row – signal recorded in the DLOR domain. Lower row-signal recorded in the VMNR domain. **E.** Six examples of the mean post stimulus histogram (PSTH) response from one electrode (each from a different electrode) on the different tasks in the two sub-regions of the STN. Upper row - recordings from the DLOR domain. Lower row - recordings from the VMNR domain. Abbreviations: DLOR-Dorsolateral oscillatory region. VMNR – ventromedial non-oscillatory region.

### Signal processing and analysis

Reaction time: Reaction time is usually defined as the time from tone onset to the start of the movement, but here we refer to the time from tone onset to the actual press. Because of the rhythmic nature of the tasks and many anticipatory (pre-tone) presses (see Table 1), anticipatory press was defined as a negative reaction time (200-0ms before tone onset). Since trial duration was 1250ms, a press more than 1050ms after the tone was considered an anticipatory press of the next tone. To avoid bias caused by repeated measures (dependency between reaction time from the same patient) in the statistical comparison (Vasey et al., 1987), the average reaction time was calculated for each session (i.e. a series of 120 tones). Responses in the All-Go task and following the go cue on the Go-NoGo task were classified as correct responses whereas the responses subsequent to the no-go cue of the Go-NoGo task were classified as commission errors (incorrect responses).

**Table 1.** Demographics, Number of Participants and Recording Sites and Behavioral Results on each o Task. Demographics: demographics and the clinical state for each participant were collected and the average and standard deviation were calculated for the participants within each task paradigm group. A one-way ANOVA tested for the effects of the demographics and clinical state between groups. Some of the clinical state data was missing for several patients who were evaluated by other medical centers. Number of recording sites and participants: each patient participated in only one task paradigm but repeated the task several times while different sites within the STN were recorded. In most cases, in each session there were two parallel recording electrodes and each electrode was considered as one recording site. Behavioral results: the behavioral results (press rate, correct / incorrect responses and reaction time) refer to the total STN recording sites. No significant changes were found between the behavioral results recorded in the DLOR and VMNR. The p values for the behavioral results are for two sample t-tests. The variables for the two-sample t-test were the averages of each recording site (i.e. average press rate, reaction time or pre-tone press rate for each recording site). Abbreviations: UPDRS-III- Unified parkinsonian rating scale, part III; LED-Levodopa equivalent dosage; ACE- Addenbrooke’s cognitive examination; FAB-Frontal assessment battery.

|  | None-Go | All-Go | Go-NoGo | p value |
| --- | --- | --- | --- | --- |
| **Demographics and clinical state** |  |  |  |  |
| Number of participants | 7 | 7 | 29 |  |
| Males:Females | 4:3 | 5:2 | 16:13 |  |
| Age (years)<br>(mean ± sd) | 67 ± 6.27 | 62.14 ± 9.08 | 62.21 ± 8.33 | $F(2,40) = 1.55, p > .05$ |
| Disease Duration (years)<br>(mean ± sd) | 14 ± 6.1 | 9.33 ± 4.03 | 8 ± 3.37 | $F(2,37) = 4.37, p < .05$ |
| UPDRS III Off Medications<br>(mean ± sd) | 43 ± 19.6 | 37.8 ± 12.79 | 43.17 ± 15.17 | $F(2,34) = .66, p > .05$ |
| UPRDS III On Medications<br>(mean ± sd) | 16.6 ± 7 | 11.2 ± 1.64 | 14.96 ± 11.44 | $F(2,34) = .36, p > .05$ |
| LED (mg/day)<br>(mean ± sd) | 1014 ± 348 | 782 ± 274 | 703 ± 453 | $F(2,38) = .76, p > .05$ |
| ACE<br>(mean ± sd) | 80.17 ± 9.43 | 81.71 ± 13.16 | 84.09 ± 10.58 | $F(2,36) = .64, p > .05$ |
| FAB<br>(mean ± sd) | 13.75 ± 2.06 | 13.57 ± 3.15 | 15.48 ± 3.41 | $F(2,34) = 1.37, p > .05$ |
| **Participants and numbers of recording sites** |  |  |  |  |
| Dorso-lateral oscillatory region (DLOR) |  |  |  |  |
| Participants no. | 5 | 5 | 23 | - |
| Recording site no. | 14 | 13 | 39 | - |
| Ventro-medial non-oscillatory region (VMNR) |  |  |  |  |
| Participants no. | 45 | 5 | 26 | - |
| Recording site no. | 14 | 13 | 80 | - |
| Total STN |  |  |  |  |
| Participants no. | 7 | 7 | 29 | - |
| Recording site no. | 28 | 26 | 119 | - |
| Recording site per participant<br>(mean ± sd) | 4 ± 1.9 | 3.71 ± 2.47 | 4.0 ± 2.17 | - |
| **Behavioral results** |  |  |  |  |
| Press rate (mean ± sd) | | | | |
| Frequent tone | - | 89 ± 14% | 94 ± 9% | $p < .05$ |
| Deviant tone | - | 85 ± 21% | 37 ± 25% | $p < .001$ |
| Reaction time (seconds, mean ± sd) | | | | |
| Frequent tone | - | 0.291 ± 0.07 | 0.413 ± 0.16 | $p < .05$ |
| Deviant tone | - | 0.367 ± 0.13 | 0.307 ± 0.19 | $p > .05$ |
| Pre-tone press rate |  |  |  |  |
| Frequent tone | - | 10.6 ± 11.2% | 3.6 ± 5.4% | $p < .001$ |
| Deviant tone | - | 8.8 ± 10.8% | 9.9 ± 21% | $p > .05$ |

Peri-stimulus histogram (PSTH): In each recording site the signal was divided into traces from 500ms before to 1250ms after the tone or press time. The root mean square (RMS) of the signal was computed in windows of 100ms, with an overlap of 50% between windows, resulting in a time resolution of 50ms bins. After calculating the root mean square values of all windows each trace was normalized by a modified Z-score. The modified Z-score was based on the median and MAD (median absolute deviation) corrected by 1.4826 (a scaling factor to equal the standard deviation (Rousseeuw et al., 1993)). The modified Z-score was chosen because it is less affected by extreme values than the Z-score. The data consisted of relatively short trials and long responses that sometimes lasted most of the duration of the trial; therefore, we chose the more resilient modified Z-score transformation. Trials were aligned to tone onset and to press time and categorized into frequent or deviant tones. The mean of all trials (modified Z-scores of the root mean square as a function of time) in each recording site was calculated before averaging all recording sites for the same sub-area of the STN. Different analysis methods (median calculated on all recording sites and mean and median calculated for all traces) yielded similar results. In order to measure the evoked response in the different categories for statistical comparison, we defined the magnitude of each response as the difference in the evoked response between the maximal peak time and the minimal trough time of the average evoked response. The maximal and minimal point were determined according to the average response within each category (see Fig. 6A, B, darker dots that mark the maximal and minimal points).

Artifact removal: artifacts in the raw data were detected by the automatic rejection criterion of an absolute amplitude exceeding 20 times standard deviation (SD). Epochs with artifacts were removed from the database and analysis. Speaker’s echo of the auditory signal picked up by the recording electrodes was filtered using a narrow filter at the auditory signal’s pitch frequencies and its harmonics and verified by a human expert (OM). Trials with Z-scored PSTH responses exceeding 6 times the standard deviation of the signal were excluded from the analysis.

### Statistics and software

Patients’ demographics in all three groups were compared by one-way ANOVAs. For each session of the All-Go and Go-NoGo tasks, we calculated the average reaction time, the average press rate and the average early-press rate for frequent and deviant tones. We tested differences in reaction time by a mixed design ANOVA with the task as the between- subject factor and the tone as the within- subject factor. In the case of a significant effect, we used post-hoc simple main effect analysis with a Bonferroni correction for multiple comparisons. In addition, we tested for difference in the early press rate between the All-Go and Go-NoGo task using two sample t-tests and repeated the test for frequent and deviant tones.

#### Neuronal data

We measured two parameters of the evoked response: the average amplitude and the latency of the peak. We first conducted a paired t-test to compare the response amplitude for the frequent and deviant tones in each of the four sub regions tested (left/right, DLOR/VMNR). Next, a two-sample t-test was conducted to compare response amplitude and latency in the left and right recording sites in both DLOR and VMNR sub-domains. We used a two-sample t-test since 25/43 and 14/43 of the patients did not have both left and right recording sites or both DLOR and VMNR recording sites, respectively. Because of the small sample size, after verifying that there were no significant differences between left and right, the data from bilateral sites were pooled. We then conducted again a paired t-test to compare the amplitudes of the evoked response to frequent and deviant tones in the pooled ventral and dorsal locations for each task. The task effect on the amplitude of neural evoked responses was compared using a one-way ANOVA test (3 groups: None-Go, All-Go, Go-NoGo) and repeated for each tone (frequent, deviant) and for each subdomain (DLOR, VMNR). In the case of significant effects, we used post-hoc simple main effect analysis with a Bonferroni correction for multiple comparisons. In the All-Go and Go-NoGo task where we also aligned the evoked responses to the press times, we compared the task effect on the response amplitude for each of the tones (frequent/deviant, two sample t-test), and the tone effect on each of the tasks (All-Go, Go-NoGo, paired sample t-test). In order to test whether the response was more affected by the tone or by press execution we compared the response amplitude of the evoked responses aligned to the tone to those aligned to the press (paired sample t-test).

To further compare the whole pattern of All-Go and Go-NoGo responses, a paired sample t-test on each of the bins in the trial (50ms bins) was conducted with FDR correction, and repeated for each tone (frequent/deviant) and for each alignment (tone/press).

In the Go-NoGo task, we further divided the response to the deviant tone into successful and unsuccessful trials. To test for significant changes in the neuronal response between the successful and unsuccessful stop trials we used a paired sample t-test with FDR correction for all the PSTH bins (50ms windows). All the analyses were 2 tailed with a significance level of a= .05. Data were processed and analyzed using Matlab 2016b (Mathworks, Inc., Natick, MA) and using SPSS 24.0 (IBM SPSS Statistics for Windows Armonk, NY: IBM Corp).

### Data availability

Data will be available at http://basalganglia.huji.ac.il/links.html

## Results

Neuronal activity was recorded through microelectrodes during DBS surgery targeting the STN. Each patient engaged in one of the following tasks: 7 patients participated in the None-Go tasks (28 recording sites), 7 patients participated in the All-Go task (26 recording sites) and 29 patients participated in the Go-NoGo task (119 recording sites). Demographics, clinical assessments and active medications are listed in Table 1. No significant differences were found between the task groups except for disease duration (one-way ANOVA). The number of recording sites and behavioral results are summarized in Table 1.

### Behavioral results

The behavioral results are presented in Fig. 1. Reflecting the task instructions, the press rate following the deviant tone of the Go-NoGo task (37±25%) was lower than the press rate following the frequent tone (94±9%, *p*<.001), and lower than the press rate on the All-Go task (85±21%*, p*<.001, Fig. 1B). The early press rate following the frequent tone was significantly higher on the All-Go task than on the Go-NoGo task (10.6±11.2% and 3.6±5.4% respectively, *p*<.001, Fig. 1B). A mixed design ANOVA, with task (All-Go, Go-NoGo) as the between-subject, and tone as the within- subject showed no significant main effect for tone type (*F*(1,136)=2.55,*p*>.05) or task (*F*(1,136)=1.49 *p*>.05) but a significant interaction between task context and tone type (*F*(1,136)=19.6 *p*<.001). Post Hoc simple main effects indicated that the reaction times after frequent tones were significantly faster on the All-Go task (.29±.07s) than on the Go-NoGo task (.41±.16s, *p*<.001) but not on the deviant tone task (.37±.13 vs. .31±.19, *p*>.05, Fig. 1C). This result reflects the repetitive nature of the All-Go task. Post Hoc simple main effects revealed that on the Go-NoGo task, the reaction times after the deviant tone (.31±.19) were faster than the reaction times after the frequent tone (.41±.16, *p*<.001, Fig. 1D). No significant effect for tone was found on the All-Go task. This was expected since the reaction time after the deviant tone (no-go cues) in the Go-NoGo task only represents commission errors (incorrect responses).

### The STN subdomains respond to the All-Go task (motor activation) with different timing

The recording locations in the STN subdomains, an example of raw recordings (high-pass filtered) and typical responses to each of the tasks are presented in Fig. 2. The average responses on the three tasks in each of the four STN subdomains (left/right, DLOR/VMNR) are presented in Fig. 3 – Fig. 5. The average evoked response to the auditory cues in the None-Go task seems to be small, as presented in Fig. 3. To estimate the magnitude of the evoked response, we defined the amplitude as the difference between the time points of the peak and the trough of the average evoked response. No significant difference between the frequent and deviant tone was detected in the DLOR and VMNR (*p*>.05; paired t-test).

**Figure 3.**
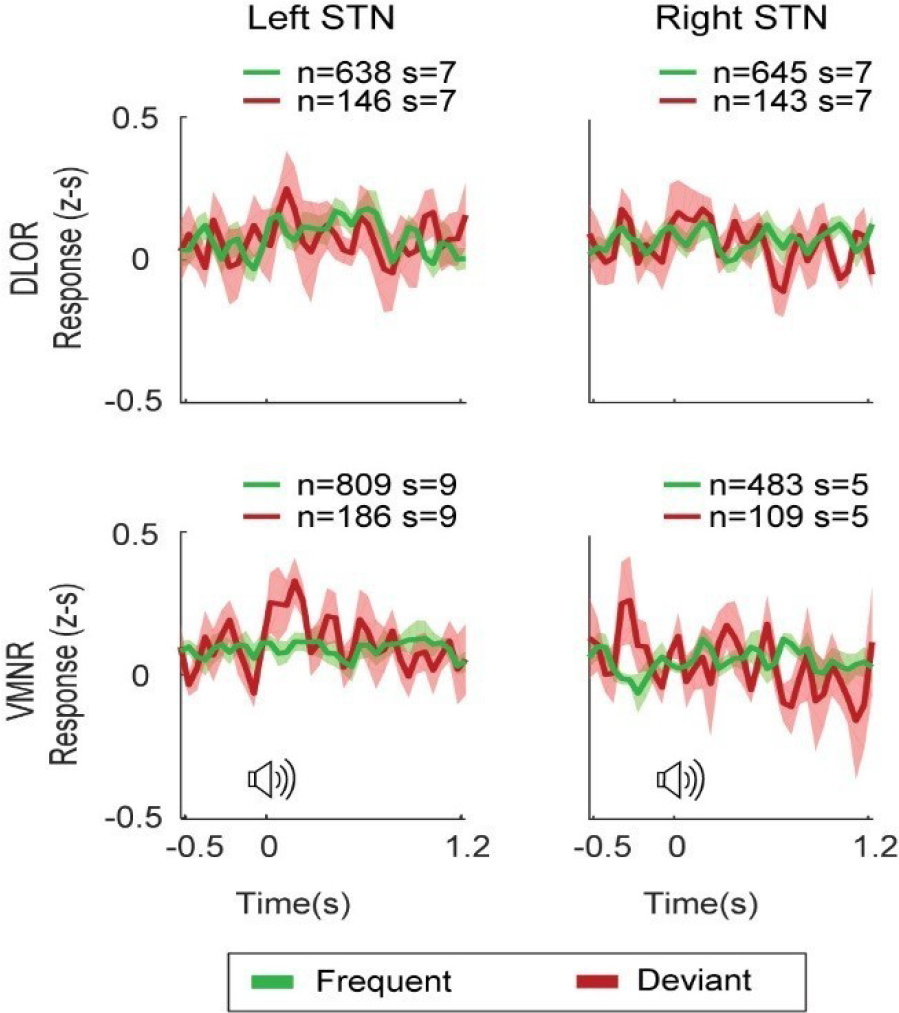
Small Response in the None-Go Task. Mean Z-scored (z-s) post stimulus histogram (PSTH) responses of microelectrode multi-unit recordings ± standard error of mean (SEM) aligned to tone in the left and right DLOR (upper row) and VMNR (lower row). n, number of trials; s, number of recording sites.

In the All-Go task, the evoked responses were seemed to be larger for both the frequent and deviant tones (Fig. 4). No significant difference was found between the evoked response amplitudes to frequent and deviant tones on the All-Go task in any of the sites except the right DLOR (*p*>.05; paired t-test). However, the timing of the average evoked response peaks to the frequent and deviant tones in the All-Go task were earlier in the VMNR than in the DLOR: the average peak response times were in the range of -0.2s to 0.2s in the VMNR (for left/eight, frequent/deviant) and in the range of 0.5s to 0.8s in the DLOR (for left/right, frequent/deviant). A two-sample t-test on the evoked response peak latencies of each recording site in the DLOR (left and right) vs. latencies of the VMNR recording sites was significantly different for the frequent tone (*p*<.01) and not significant for the deviant tone (*p*=.057).

**Figure 4.**
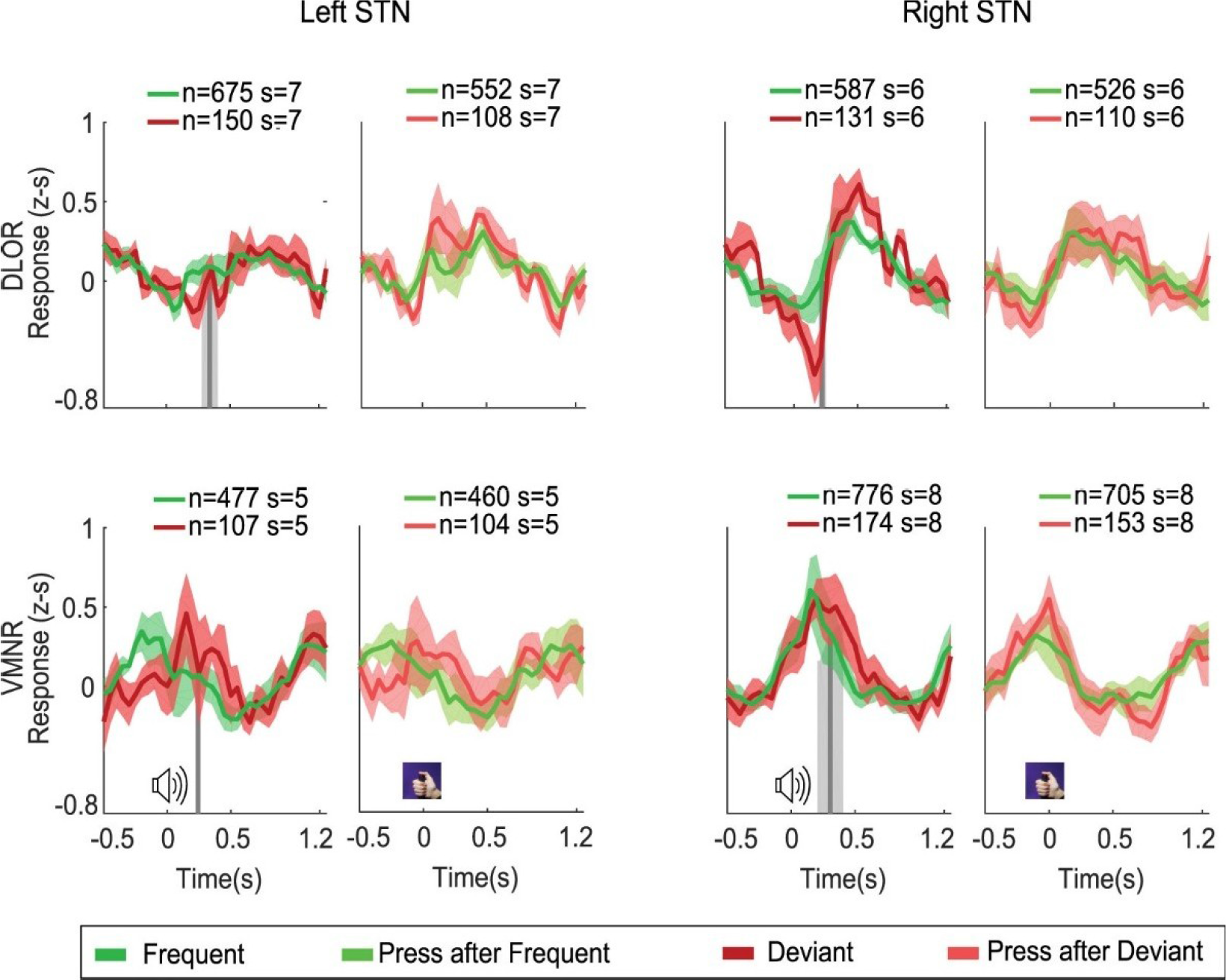
Similar Responses to the Frequent and Deviant Tones in the All-Go Task. Mean Z-scored post stimulus histogram (PSTH) responses of microelectrode multi-unit recordings ± standard error of mean (SEM, shadow) aligned to the time of the tone (left) and the time of the press (right) in the left and right STN, DLOR (upper row) and VMNR (lower row). Gray vertical lines are the mean reaction times of the frequent tone with its standard deviation (shaded). Green and red lines are the frequent and deviant tones, respectively aligned to the time of the tone; light red and light green lines are the frequent and deviant tones respectively aligned to the press. n, number of trials; s, number of recording sites.

### Differential responses to the Go vs NoGo cues in the limbic-associative domain of the STN

In contrast to the All-Go task, the Go-NoGo task was characterized by disparate responses to the frequent tone (go signal) and deviant tone (no-go cue) in both the VMNR and the DLOR (Fig. 5). In the VMNR, the amplitudes of the evoked responses to the frequent tone were significantly smaller than the evoked responses to the deviant tone (*p*<.01, *p*<.05, left and right, respectively; paired t-test). As in the All-Go task, the average evoked response peaks of the frequent and deviant tones in the Go-NoGo task was earlier in the VMNR (in the range of 0.1s to 0.3s) than in the DLOR (in the range of 0.4s to 1s). However, a two-sample t-test on the latencies of the evoked response peaks indicated no significant differences (*p*>.05). In the DLOR, the average evoked responses were larger in alignment with the press than in alignment with the tone (significant for left DLOR, non-significant for right DLOR, *p*<.05; paired t-test). In the VMNR, the evoked responses seemed to be more locked to the tone than to the press (Fig. 5, lower row).

**Figure 5.**
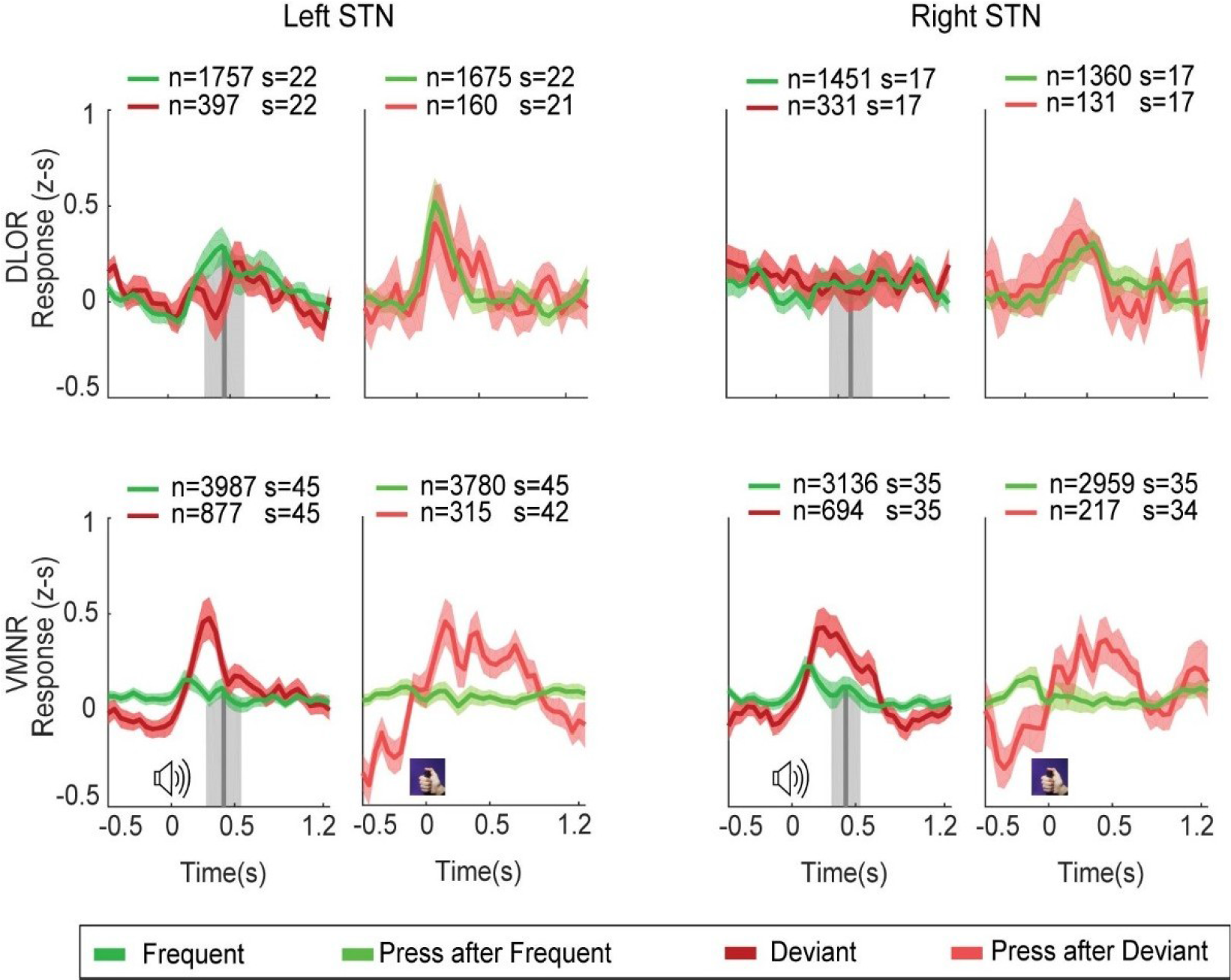
Smaller Response to the Frequent Tone in the Go-NoGo Task, in the Ventro-medial STN. Mean Z-scored post stimulus histogram (PSTH) responses of microelectrode recordings± standard error of mean (SEM, shadow) aligned to the tone (left) and to the press (right) in the left and right STN, DLOR (upper row) and VMNR (lower row). Gray vertical lines are the mean reaction times of the frequent tone with its standard deviation (shaded). Green and red lines are the frequent and deviant tones, respectively aligned to the time of the tone; light red and light green lines are the frequent and deviant tones, respectively aligned to the press. n, number of trials; s, number of recording sites.

### Evoked response to a Go cue decreases in the context of movement inhibition in the limbic-associative domain of the STN

Although the patients pressed the button with their right thumb or index finger, evoked responses to movement were observed in both the left and right STN. No statistical difference was found between the left and right STN amplitudes or latencies on overall, except three cases (the amplitude of the deviant tone of the All-Go task in the DLOR (*p*<.05, two sample t-test), the latencies of the deviant tone on the All-Go task in the DLOR and on the Go-NoGo task in the VMNR (*p*<.05, two sample t-test)). In order to increase statistical strength and to compare the evoked responses between the different tasks, we pooled the left and right STN recording sites (Fig. 6). We repeated the paired t-test analysis to test for differences between the frequent and deviant tones’ evoked response amplitudes that was conducted above in the four STN sub-regions for each task. The VMNR response to the deviant tone in the Go-NoGo task and in the None-Go task was larger than its response to the frequent tone (*p* <.001, *p*<.05, respectively) (Fig. 6A, C). All other comparisons were non-significant.

**Figure 6.**
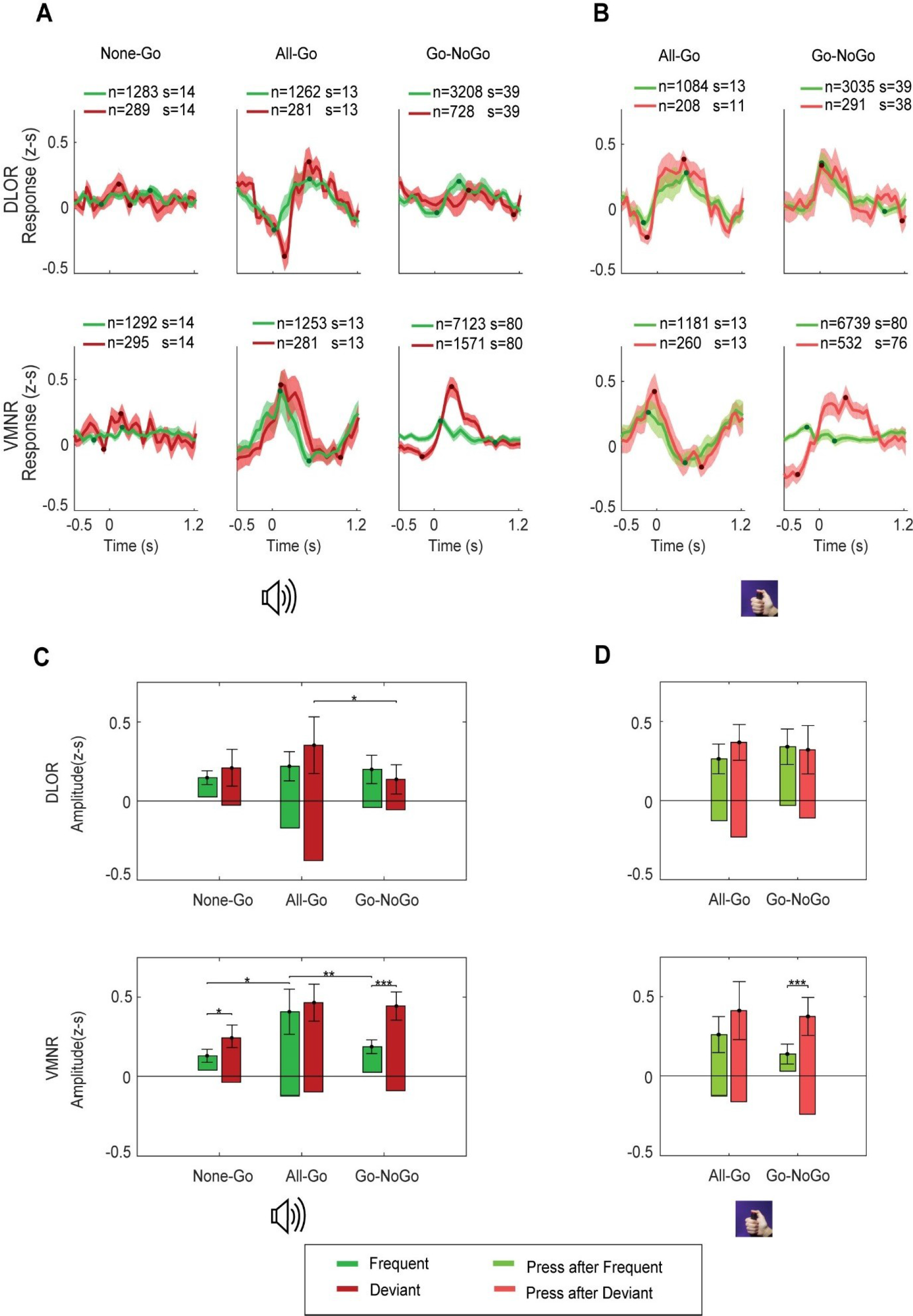
Responses to the Frequent Tone in the VMNR Decrease in the Context of Movement Inhibition. **A.** Average PSTH (post stimulus histogram) response to the tone in the DLOR (upper row) and VMNR (lower row) from all recording sites (left and right STN together) on the three tasks, aligned to tone. Shaded areas represent the standard error of the mean. Darker dots represent the time used for the amplitude measure. N, number of trials; s, number of recording sites. **B.** Average PSTH (post stimulus histogram) response aligned to the press. Same conventions as in A. **C.** Amplitudes of the average responses aligned to tone in each task in the DLOR (upper row) and the VMNR (lower row). **D.** Amplitudes of average frequent and deviant responses aligned to press for the All-Go and Go-NoGo tasks in the DLOR (upper row) and VMNR (lower row). The amplitude of the average signal was defined as the difference between the maximal peak and the minimal trough of the evoked response. Statistical difference between the tasks was calculated by a one-way ANOVA on the amplitudes of each response at the times of the average responses’ maximal and minimal points. Post Hoc statistical analysis was calculated with a Bonferroni correction. Statistical differences between the tones on each task were calculated by a paired t-test.

To test the effect of task context on STN evoked responses, we conducted a one-way ANOVA on the amplitude of the evoked response to the tone, with the task context as a parameter (None-Go, All-Go, Go-NoGo). We repeated the analysis for the frequent and deviant tones and for the VMNR and DLOR sub-regions of the STN. The VMNR response to the frequent tone was modulated by task context (*F*(2,104)=5.56, *p*<.01). Post-hoc pairwise comparisons revealed that the evoked responses to the frequent tone were lower in the None-Go and Go-NoGo tasks than in the All-Go task (*p*<.05 None-Go, *p*<.01 Go-NoGo). Surprisingly, there was no effect for task context on the responses to the deviant tone (*F*(2,104)=2.7 *p* >.05). In the DLOR, task context did not modulate responses to the frequent tone but did affect responses to the deviant tone (*F*(2,63)=1.08, *p*>.05, *F*(2,63)=4.37, *p*<.05, respectively). Post-hoc pairwise comparisons for the deviant tone showed larger responses on the All-Go than on the Go-NoGo task, which was expected given the behavioral difference seen above; i.e., 37% of the tones were followed by a press on the Go-NoGo task compared to 85% on the All-Go task (Fig. 1B). No other pairwise comparisons were significant (*p*>.05).

### The smaller amplitude after the Go cue on the Go-NoGo task is mainly due to a deficiency in the amplitude’s negative component

To further compare the differences between the response pattern on the All-Go and Go-NoGo tasks, we superimposed the All-Go and the Go-NoGo responses with a normalization to the time before tone or press (Fig. 7). In the VMNR, the All-Go task evoked response aligned with the press had a large negative component (i.e., a reduction in neuronal activity) at 400-600ms after the press on the frequent tone trials and 250-850ms after the tone on the deviant tone trials that was not observed on the Go-NoGo task (marked by gray areas, tested for significance level of *p*<.05, paired t-test after FDR correction). In the DLOR, the negative component of the All-Go response was more prominent when the responses were aligned to the tone, and significantly different from the Go-NoGo for the deviant tone at 150-250ms after tone onset.

**Figure 7.**
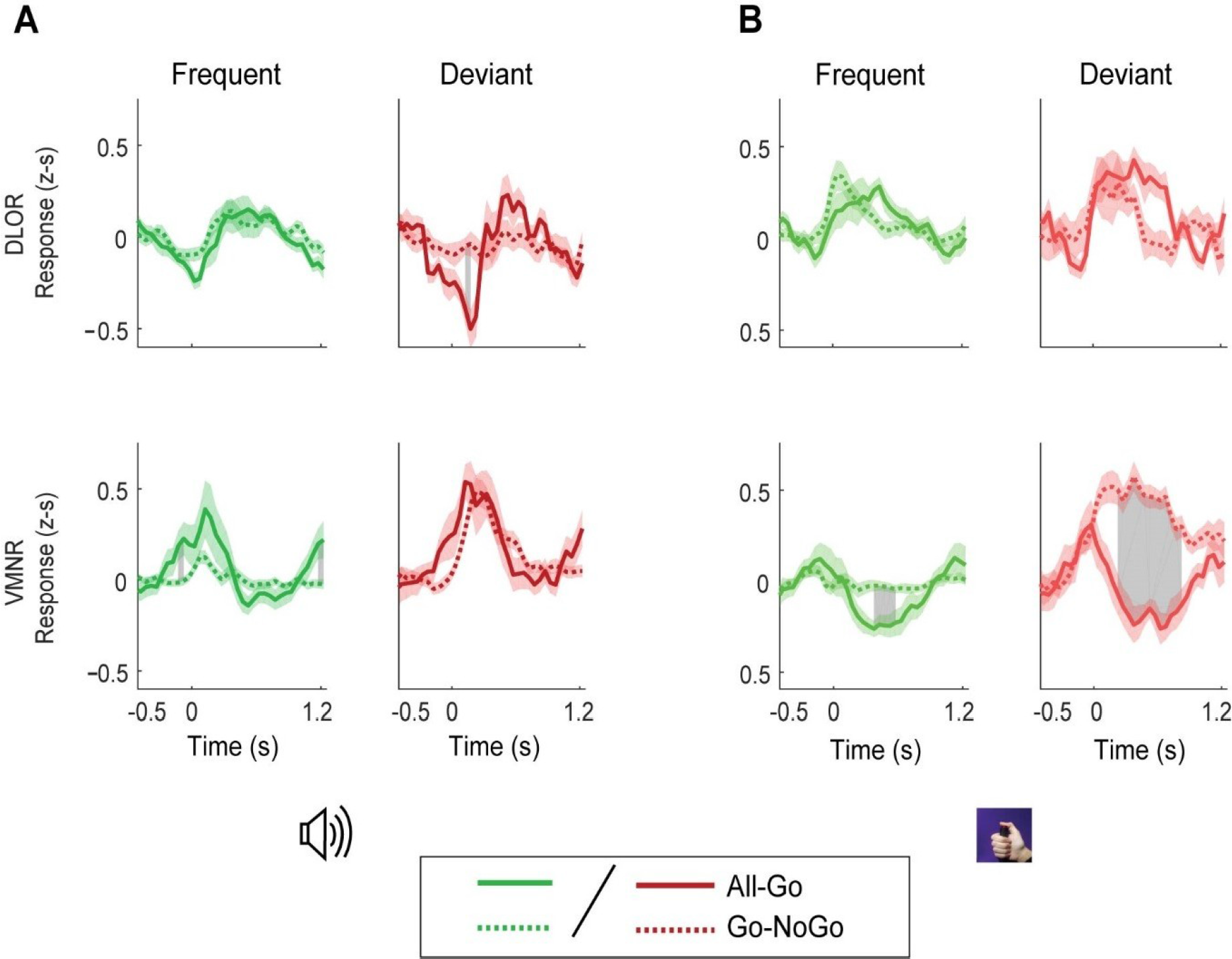
Disappearance of the Responses’ Negative Phase in the Context of Movement Inhibition. **A.** Mean PSTH response of all recording sites aligned to the tone in the All-Go (solid line) and Go-NoGo (broken line) for the frequent (green) and deviant (red) tones. The responses were normalized by subtraction of the mean activity at the time before tone onset (−500:−100ms). **B.** Mean PSTH response of all recording sites aligned to the press. Same convention as in A. Gray area indicates time bins in which the difference between the All-Go and Go-NoGo tasks was significant (two-sample t-test *p*<0.05 after FDR correction).

### Commission Error responses are associated with lower neuronal activity before tone onset

Analysis of the response to the no-go cue (deviant tone) on the Go-NoGo task indicated a change in neuronal activity between the correct rejection (omission) and commission error trials (Fig. 8). In the VMNR, the neuronal activity on trials with commission errors was significantly lower than the neuronal activity on trials with correct rejection at 500-100ms preceding the tone (tested for significance level of *p*<.05, paired t-test after FDR correction). The commission error trials displayed a significantly elongated phase of increased neuronal activity 500-1000ms after the tone.

**Figure 8.**
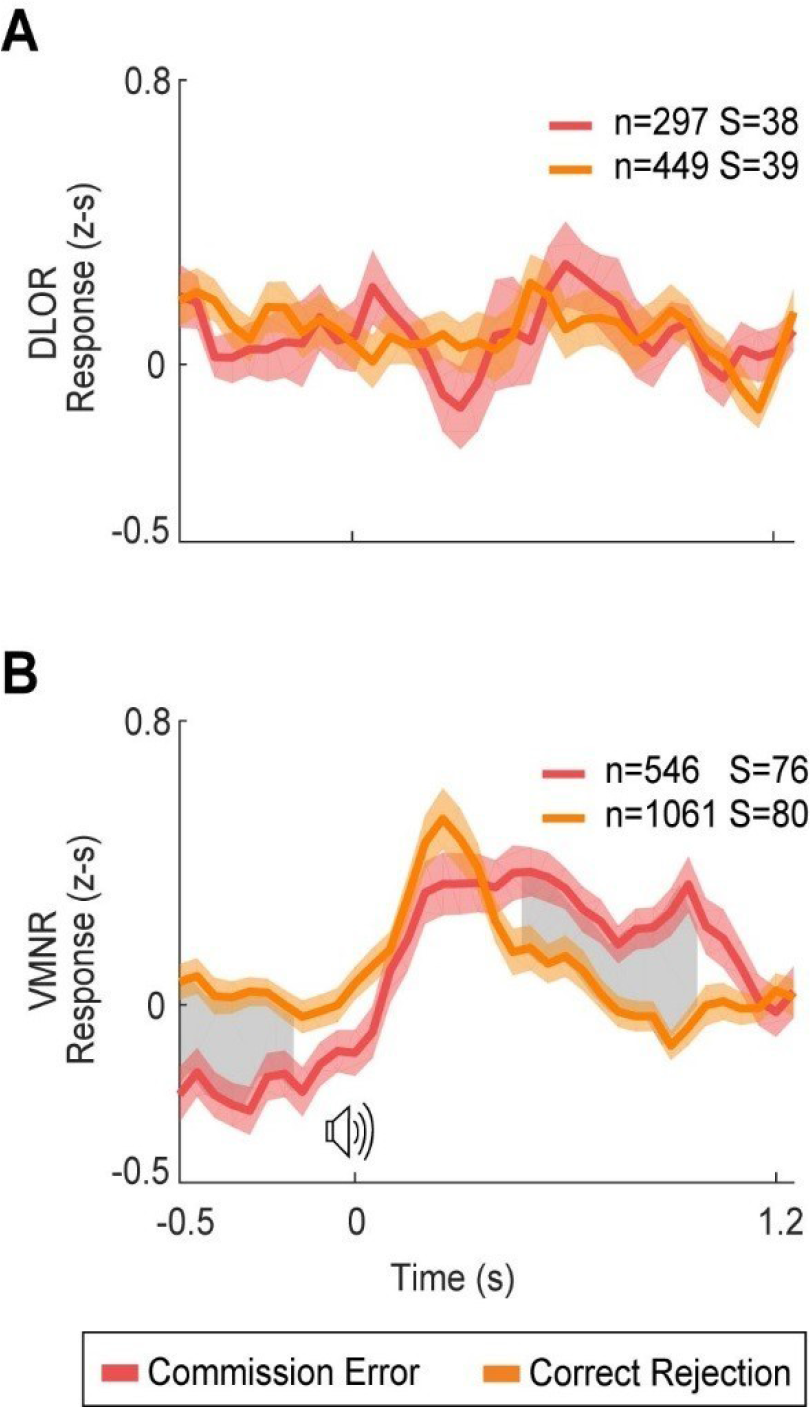
Differences between Correct Rejection and Commission Error Responses in the Go-NoGo task. Mean PSTH responses of all recording sites for trials of correct rejection (orange) and commission errors (pink) in response to the No-Go signal on the Go-NoGo task in the DLOR (A) and VMNR (B). Shadows represent the standard error of the mean. Gray area indicates time bins in which the difference between the commission error and the correct rejection responses was significant (paired-sample t-test *p*<0.05 after FDR correction). n, number of trials; s, number of recording sites.

## Discussion

The findings revealed significant changes in human STN multiunit activity between the three levels of an oddball task. The ventro-medial STN adapts the response to movement inhibition context by selectively decreasing neuronal activity. Here, although the response to the deviant tone remained the same in the context of motor facilitation (All-Go task) and motor inhibition (Go-NoGo task), the response to the frequent tone (go cue) decreased in the context of motor inhibition (Go-NoGo task). The decrease in amplitude of the evoked response on the Go-NoGo task was mainly in the negative component of the response. In addition, the commission error responses (incorrect responses to an inhibitory signal) had a larger negative component before the tone and an elongated positive phase after the tone compared to the omission responses (correct responses to an inhibitory signal).

### The ventromedial STN corresponds to movement planning while the dorsolateral STN corresponds to movement execution

Analysis of both the All-Go and Go-NoGo tasks revealed different responses in the VMNR and DLOR that probably represent their different roles in movement planning and movement execution. The current data support the view that VMNR activity is related to movement planning whereas DLOR activity is related to movement execution. First, the VMNR responses preceded the DLOR responses. For example, the peak of the evoked response to frequent tone, that represents the majority of the presses, was earlier in the VMNR than in the DLOR, both on the All-Go and Go-NoGo tasks (0.15s vs. 0.55s and 0.1s vs. 0.4s after the tone, respectively) and the evoked response started before the press time in the VMNR, both on the All-Go and Go-NoGo tasks (0.1s and 0.15s before the press, respectively, Fig. 6B, bottom right row). Second, the VMNR responses are not correlated to execution of movement. For example, large evoked response amplitudes were observed to the deviant tone in the Go-NoGo task, which on most trials, was not followed by movement. Third, DLOR responses were larger when aligned to the press than aligned to the tone (Fig. 4, Fig. 5, upper row). Finally, the peak of the DLOR responses were usually around the mean reaction time (most prominently in the left DLOR, Go-NoGo task, aligned to tone, see Fig. 5, upper row, left). The bilateral activation in response to the right thumb press might hint that the role of the STN has more to do with global movement regulation and not the execution of specific muscles.

For the first time this study reports the involvement of the ventral STN (non-motor domain) in movement planning that is not restricted to movement inhibition. These results are in line with previous findings in LFP. Motor execution has been associated with dorsal STN located DBS contacts exhibiting high beta power (Kuhn et al., 2004, Androulidakis et al., 2008, Zaidel et al., 2010, Greenhouse et al., 2011). Movement inhibition that is part of the movement planning process has been associated with ventral-STN located DBS contacts (Hershey et al., 2010, Alegre et al., 2013) and also in single units recorded from the ventral STN areas of OCD patients (Bastin et al., 2014).

### Maximal neuronal response decreases in the context of motor inhibition

The None-Go task is a passive auditory discrimination process; i.e., the detection of a change in the auditory pitch similar to the mismatch negativity test. The None-Go task elicited small STN evoked responses but with a significant difference between the frequent and deviant tones in the VMNR (*p*<.05). This might contribute small effect to the STN responses on the two other tasks (All-Go and Go-NoGo tasks).

Surprisingly, we found that the STN evoked responses to the frequent tone in the Go-NoGo task were lower than the evoked responses in the All-Go task. Previous studies that have examined inhibitory paradigms reported a stronger evoked response to an inhibitory signal and thus suggested a mechanism of increased activation of the STN to the inhibitory signal (Aron and Poldrack, 2006, Isoda et al., 2008, Benis et al., 2016, Wessel et al., 2016a). Our results suggest that the response to the inhibitory cue does not increase; rather the response to the go cue decreases. Below we discuss three possible, non-mutually exclusive explanations for the decreased STN response in the context of motor inhibition. The first has to do with resource allocation in the STN subsequent to an increased cognitive load. The second focuses on the modulation of STN neuronal activity as a mechanism of facilitation and inhibition of movement. The third relates to the process of error monitoring in the STN.

#### 1. Allocation of neuronal resources to the different cues compensates for the STN capacity constraints

The decreased STN responses in the motor inhibition context (the most complex of the three tasks in this study) may represents a limitation in the computational capacity of STN neurons. We suggest that this limited capacity may result in a selective integration of data involving concentration on relevant stimuli and a filtering out of non-relevant stimuli. If a motor task also requires preparation for possible movement inhibition, the neuronal capacity reserve decreases, and the allocation of resources to the frequent and deviant tones reflects the main aim of the task. In the simpler task (All-Go) the neuronal responses to the frequent tone are similar to the deviant tone in the VMNR. Since the aim of the All-Go task is motor planning and execution, task completion does not require discrimination between the frequent and deviant tones. In the more complex task (Go-NoGo) the neuronal responses in the VMNR (which is associated with movement planning) were found to be smaller to the frequent tone than to the deviant tone (the amplitude/ response ratio was 0.3:1, for the frequent vs. deviant tone respectively). The major aim of the Go-NoGo task is inhibition of action (press) upon the deviant tone (no-go cue). Thus, more neuronal resources should be allocated to the no-go cue. Since increasing the response to the no-go cue is not possible due to the limited neuronal capacity, the discrimination between the go and no-go cues takes the form of a reduction of the responses to the go cue. The supposition that the STN can prioritize the process that is the most relevant to the task is in line with previous reports (Baunez et al., 2001, Wessel et al., 2016a).

#### 2. Preparation for movement inhibition decreases the fluctuations in STN activity during movement

In the classical model of the basal ganglia, the role of the STN is to provide ongoing continuous (tonic) inhibition (“brakes”) on movement execution. The high spontaneous STN firing rate represents the baseline tonic inhibition and the decreased STN firing rate represents a release of this tonic inhibition. In the current study, the movement in the Go-NoGo task is more restrained due to the ongoing preparation for the no-go cue, whereas in the All-Go task the movement is more free and rhythmic (i.e. uninterrupted) due to the fixed inter-tone interval that encourage movement anticipation. In the All-Go task, the repetitive nature of the movement is reflected behaviorally by a shorter reaction time and an increased percentage of early presses (before tone onset). In line with these behavioral changes, in the All-Go task there is a larger negative component (i.e. lower neuronal activity) that precedes the evoked response, which may represent a release of tonic inhibition (see Fig. 6A, and Fig. 7A,). In the Go-NoGo task, the absence of a negative component may reflect ongoing tonic inhibition. Besides the absence of the negative component, the evoked responses are also narrower in the context of movement inhibition than the same evoked responses within the context of movement facilitation (see Fig. 7B, upper row). This shorter response might represent the readiness of the STN for the anticipated inhibitory signal. Another supporting finding for decreased fluctuations of neuronal activity as a mechanism that facilitates movement inhibition is the correlation between the level of neuronal activity before the tone and the ability to inhibit movement in the Go-NoGo task. Decreased neuronal activity before the inhibitory signal in the Go-NoGo test is correlated with the inability to inhibit the movement; i.e., commission error, whereas higher neuronal activity before the inhibitory signal is correlated with the inhibition of movement; i.e., correct rejection (see Fig. 8).

Our claim that the level of modulation in STN activity corresponds to the level of action control (i.e. the context of movement facilitation vs. the context of movement inhibition) is supported by recent studies. Greenhouse et al. (2015) reported that the level of motor-evoked potential inhibition during response preparation was sensitive to response complexity. Fischer et al. (2016) described a cortical mechanism of decreased amplitude in the movement response when adding anticipation of movement inhibition to regular repetitive tapping. They reported that successful motor inhibition was associated with increased beta power activity in the parietal region EEG prior to the inhibitory signal. Benis et al. (2014) reported that unsuccessful motor inhibition trials had relatively lower beta-band (13-35Hz) LFP activity in the STN after cue onset. The level of STN LFP beta power modulation during movement was also reported to be correlated with motor performance (Androulidakis et al., 2007, Tan et al., 2015, Fischer et al., 2017b). However, these reports are based on STN or cortical LFP, whereas the current results draw on the rate and pattern of multiunit activity.

#### 3. Error monitoring in the STN in the context of movement inhibition

The basal ganglia play a major role in reinforcement learning by monitoring the error between the prediction and the actual outcome. Animal studies suggest that dopaminergic neurons fire briefly around the prediction and reward times and that the magnitude of their firing rates encodes the difference (error) between the prediction and the actual outcome (Wise et al., 1989, Schultz et al., 1992, Pizzagalli et al., 2008, Joshua et al., 2009b). More specifically, dopaminergic neurons play a role in error monitoring of movement feedback (Morris et al., 2006, Joshua et al., 2009a). Although the STN receives only a small fraction of dopaminergic projections compared to the striatum (Rommelfanger et al., 2010), several studies in rats and recently also in human subjects have reported that the STN is also involved in error monitoring (Lardeux et al., 2009, Baunez et al., 2011, Lardeux et al., 2013, Bastin et al., 2014, Tan et al., 2014, Breysse et al., 2015). Analysis of changes in post-press neuronal activity that represents the error monitoring phase in the STN hinted here that the DLOR post-press period may represent the motor execution feedback. The post-press period is characterized by increased neuronal activity both in the context of movement facilitation (All-Go task) and movement inhibition (Go-NoGo task, see Fig. 6B and Fig. 7B, upper rows). In contrast to the DLOR, the post-press feedback period in the VMNR depends on the movement context. In the context of movement facilitation (All-Go task) the feedback phase is characterized by decreased neuronal activity. However, in the context of movement inhibition (Go-NoGo task) the feedback period may also indicate whether the press was correct or not (error monitoring). In the case of presses after the no-go cue, the press is incorrect, thus leading to an error feedback such that the post-press period is characterized by increased activity (Fig. 6B lower rows). Although there are no data on alignment to press after a correct rejection response (because there is no press) the response aligned to tone shows an elongated increased amplitude in commission error responses compared to correct rejection responses (Fig. 8 B). This differential response may represent the level of error (“oops response” (Lardeux et al., 2009)). The feedback to the go-cue in the Go-NoGo task is on the one hand motor feedback or preparation for the next press (reflected in decreasing neuronal activity, as in the All-Go context) and on the other hand error monitoring feedback (whether the press was correct or not) mediated by an increase in neuronal activity. Hence, the net response to the go cue in the Go-NoGo task is no change in neuronal activity.

### Study limitations

The most obvious limitation of this study is that electrophysiological investigations in Parkinson’s disease patients cannot necessarily be generalized to healthy subjects. Besides the motor symptoms, Parkinson’s disease patients exhibit a decline in response inhibition and other deficits that are related to the tasks administered here such as attention shift, error monitoring, the ability to learn from negative decision outcomes, and multitasking (Witt et al., 2004, Frank et al., 2007, Castner et al., 2008, Obeso et al., 2013, Muralidharan et al., 2016). STN electrophysiology may be influenced by the motor, emotional and cognitive symptoms of Parkinson’s disease (Cassidy et al., 2002, Levy et al., 2002, Kuhn et al., 2004, Priori et al., 2004, Eitan et al., 2013, Rappel et al., 2018). The basal STN neuronal activity in Parkinson’s disease might be higher than in healthy subjects. The STN capacity may be more limited in Parkinson’s disease, the processes of movement facilitation and inhibition may be impaired, and the error monitoring activity may be altered due to the change in dopamine levels.

To avoid learning effects on the different tasks we recorded only one task per participant in this study. One limitation of this decision is that no recordings are available from the same cell or STN domain in the same patient on different tasks. However, the advantage over many LFP studies in the STN lies in the microelectrode recordings with a discrete, small and accurate sampling area and the differentiation into sub-territories of the STN.

### Conclusion

Overall, the findings suggest that the human ventro-medial STN prepares for a possible inhibitory signal by selectively decreasing the activity modulation associated with movement. In the context of movement inhibition, the response amplitude to the release (go) cue decreases (compared to the same signal in the All-Go task) rather than increases in the response amplitude to the inhibitory (no-go) cue. This reduced response amplitude to the go signal may represent an STN mechanism for discrimination between go and no-go cues when a further increase in the response to the no-go cue is limited. The smaller amplitude of the response to the release (go) cue in the context of motor inhibition is mainly due to a smaller negative component in the evoked response. This smaller negative component may result from two possible mechanisms. First, a larger negative component (decrease in neuronal activity) represents movement facilitation whereas a smaller negative component enables a preparation for possible movement inhibition. Second, the smaller negative component may result from the error monitoring process that occurs after the press and is mediated by the level of increase in neuronal activity. Finally, the results indicate that the non-motor domain of the STN is involved in movement control and therefore should not be completely avoided during targeting of DBS contacts for the treatment of Parkinson’s disease.

## Acknowledgment

The authors acknowledge Dr. Kirk R Daffner’s contribution to the analysis and thank him for helpful advice.

